# Sex divergent behavioral responses in platform-mediated avoidance and glucocorticoid receptor blockade

**DOI:** 10.1101/2023.09.26.559122

**Authors:** Carly J. Halcomb, Trey R. Philipp, Parker S. Dhillon, J. Hunter Cox, Ricardo Aguilar-Alvarez, Samantha O. Vanderhoof, Aaron M. Jasnow

## Abstract

Women are more likely than men to develop anxiety or stress-related disorders. A core behavioral symptom of all anxiety disorders is avoidance of fear or anxiety eliciting cues. Recent rodent models of avoidance show reliable reproduction of this behavioral phenomenon in response to learned aversive associations. Here, a modified version of platform-mediated avoidance that lacked an appetitive task was utilized to investigate the learning and extinction of avoidance in male and female C57BL6/J mice. Here, we found a robust sex difference in the acquisition and extinction of platform-mediated avoidance. Across three experiments, 63.7% of female mice acquired avoidance according to our criterion, whereas 83.8% of males acquired it successfully. Of those females that acquired avoidance, they displayed persistent avoidance after extinction compared to males. Given their role in regulating stress responses and habitual behaviors, we investigated if glucocorticoid receptors (GR) mediated avoidance learning in males and females. Here we found that a subcutaneous injection (25mg/kg) of the GR antagonist, RU486 (mifepristone), significantly reduced persistent avoidance in females but did not further reduce avoidance in males after extinction. These data suggest that GR activation during avoidance learning may contribute to persistent avoidance in females that is resistant to extinction.

## 1.0 Introduction

Anxiety disorders have a lifetime prevalence of approximately 13.6-28.8% of the worldwide adult population (Michael, et al., 2007). However, women are 60% more likely than men to develop an anxiety or trauma-related disorder (McLean et al., 2011; Kessler, et al., 2012), yet it remains unclear why this difference exists. Anxiety symptomatology is often characterized by avoidance of anxiety-inducing cues (Mowrer, 1960; Hayes, et al., 1996). Animal models of avoidance have been used for decades to assess fear behavior, learning, and memory and to understand the mechanisms of anxiety and fear-related disorders as well as sex differences across various paradigms. However, most of these studies utilized inhibitory avoidance, two-way active avoidance, or one-way active avoidance tasks. Active avoidance conditioning is useful for measuring an animal’s direct response in the presence of a threat-predicting stimulus, compared to inhibitory avoidance, which relies upon the inhibition of a rodent’s natural tendency to avoid brightly lit areas. Active avoidance is useful for understanding both avoidance learning and memory and the extinction of these learned responses, to allow for better understanding of the behavioral and neural processes that regulate an animal’s response to a signaled threat. A recently developed active avoidance conditioning task is called platform-mediated avoidance (PMA). In this procedure, animals are first trained on a variable interval (VI) schedule to lever press for sucrose pellets. Then, using classical conditioning to learn an association between a neutral conditional stimulus (CS) and an aversive unconditional stimulus (US). Animals are presented with a series of CS-US pairings. Animals learn that the neutral CS (e.g., tone) predicts the aversive US (e.g., shock). However, animals also can learn to escape the US by stepping onto a nearby platform (Diehl, et al., 2019; Bravo-Rivera, et al., 2014). During this procedure, animals initially show high levels of tone-induced freezing and drastically reduced lever pressing. However, over subsequent conditioning trials, freezing decreases as animals learn to avoid the shock by stepping on the platform, and lever pressing returns to pre-conditioning levels during the inter-tone intervals (Bravo-Rivera, et al., 2014). The advantage of this design over two-way active avoidance is that there is always a safe place where the animals can avoid shock by stepping on the platform. With the addition of the appetitive task, there is a cost of avoiding the shock (i.e., loss of food reward), unlike one-way active avoidance (Diehl, et al., 2019). Following conditioning, animals can be trained to no longer fear the CS by undergoing extinction in the absence or presence of the escape platform. Extinction is widely accepted as the basis for exposure therapy (McNally, 2007), which results in learning that the CS no longer predicts the US (Bouton, et al., 2006). Following extinction, animals receive the CS alone and show a reduction in fear behavior (e.g., avoidance or freezing) due to this new learning. Overall, the PMA model is a useful behavioral paradigm for assessing avoidance responses and the extinction of learned avoidance when there is a protective mechanism present.

Although sex differences have been reported in aversive learning, the results have been somewhat equivocal regarding the presence and direction of the sex differences. For example, some studies have shown that female mice display deficits in multiple aversive conditioning paradigms (Maren, et al., 1994; Gresack, et al., 2009; Day & Stevenson, 2020). However, others have shown no sex differences (Pryce, et al., 1999) or less fear in females compared to males (Binette, et al., 2022; Baren, et al., 2009). Notably, females that acquire aversive conditioning tend to have a higher resistance to extinction of the aversive cues (Greiner, et al., 2019). Additionally, females require more safety training than males to learn conditioned inhibition (Adkins, et al., 2022). Some reports have demonstrated sex differences in fear response strategies where female mice display active avoidance over freezing (Gruene, et al., 2015; Colom-Lapetina, et al., 2019). However, it has also been reported that flight and darting responses in mice are non-associative and do not reflect associative learning (Trott, et al., 2022). Thus, how male and female rodents might differ in their fear responses across multiple behavioral outputs, including PMA, needs further attention.

Exposure to stress prior to fear extinction training has differential effects in males and females (Griener, et al., 2019; Binette, et al., 2022: Baran, et al., 2009). A key component in regulating stress is the release of glucocorticoids from the adrenal glands due to adrenocorticotropic hormone release from the anterior pituitary (McEwen, et al.,1975; Barlow, et al., 1975). Glucocorticoids bind to mineralocorticoid (MRs) and glucocorticoid receptors (GRs) to mediate physiological and behavioral responses during stress and regulate the return to homeostasis (Smith & Vale, 2006). Notably, activation or suppression of GRs can impair fear extinction (Green, et al., 2011; Camp, et al., 2012; Knox, et al., 2012), indicating a delicate balance in GR regulation of fear suppression. Similarly, corticosterone can facilitate or impede fear acquisition, expression, and extinction retention (Lesuis, et al., 2021; Thompson, et al., 2004; Skórzewska, et al., 2006; Brinks, et al., 2009). This variable behavioral response is thought to occur due to the levels of glucocorticoids present at the time of learning, where high or low levels of glucocorticoids result in learning impairments, but moderate levels facilitate adaptive synaptic plasticity (*for review see*, Sandi, 2011). The hypothalamic-pituitary-adrenal (HPA) axis is sensitive to sex hormones which may contribute to the observed sex differences in behavioral and neuroendocrine stress responses. For example, high levels of estradiol in females can result in a delayed return to baseline levels of glucocorticoids following stress, as well as an overall increase in plasma corticosterone (Carey, et al., 1995; Viau, et al., 1991). In addition, there are sex differences in GR activation (*for review see* Bourke, et al., 2012).

Females show enhanced activation of GRs compared to males within the hypothalamus following both acute and chronic stress (Zavala, et al., 2011), as well as enhanced GR expression within the dorsal hippocampus following a single prolonged stress exposure (Keller, et al., 2015). Given the complex role that GRs play in regulating aversive learning, the documented sex differences in GR activation after stress, and sex differences in the effects of stress on fear learning, dysregulation of GRs may underly the sexual dimorphism in behavioral output observed across several fear learning paradigms.

While some studies have focused on the role of glucocorticoids and GRs in inhibitory avoidance learning (Roozendaal & McGaugh, 1997; Chen, et al., 2012), there has been limited investigation of GRs in active avoidance. Measures of glucocorticoid levels across various fear learning stages (e.g., acquisition, expression, extinction) have emphasized their importance on adaptive fear learning and memory. Investigations using active avoidance are limited; therefore, the present study examined learning and extinction in male and female mice and the role that GRs play in regulating persistent avoidance. Utilizing male and female mice allows the investigation of potential GR regulation in sexually dimorphic behavioral strategies.

## 2.0 Methods

### 2.1 Animals and Housing

Sixty-eight male and 102 female 7–10-week-old C57BL/6J (Jackson Laboratories Stock #: 000664) mice were used for these studies. After exclusion due to behavioral criterion, 57 males and 65 females were used for statistical comparisons in Figures 2-4. All mice were housed on a 12:12 light:dark cycle with free access to food and water. Mice were housed in groups of 2-5 per cage. All experiments were conducted with approval from Kent State University Institutional Animal Care and Use Committee (IACUC) and the University of South Carolina IACUC and following NIH guidelines for the Care and Use of Laboratory Animals.

### 2.2 Platform-Mediated Avoidance Training

Platform-mediated avoidance learning was performed in four identical conditioning chambers (12” W x 12” D x 12”H) containing two Plexiglas walls, two aluminum sidewalls, and a stainless-steel grid-shock floor (Coulbourn Instruments, Allentown, PA). Mice were trained in these conditioning chambers with the addition of a 4” x 4”, white, square acrylic escape platform placed in the back right corner, which enabled the mice to avoid being shocked. The conditioning context consisted of grid floors, dotted background, and house light, and all chambers were cleaned with 70% ethanol. Before training, mice were pre-exposed to the conditioning context for five minutes, with the platform included. Twenty-four hours later, mice were placed back into the training context and, after a 120-second baseline, were presented with five 30-second tone-shock pairings (75 dB, 6kHz; 0.5 mA, 1s), with the shock delivered at the tone offset (zero delay), and a 182-second inter-tone interval (ITI). Mice underwent three consecutive daily training sessions, during which they had the opportunity to avoid the shocks by stepping onto the platform. All the acquisition sessions were scored for freezing, avoidance (time on the platform), and latency to mount the platform. If mice received three or more shocks on the third training session, they were excluded from further behavioral testing. Following the completion of the last training session, mice were matched based on the percent time on the platform as a measure of avoidance and were placed in extinction or context exposure groups. Twenty-four hours later, mice underwent extinction or context exposure. Extinction training occurred in the conditioning context without the platform, during which mice were presented with 30 non-reinforced tones (75db, 30 s, 6 kHz, 60ITI). Extinction sessions lasted 32 minutes. The context exposure group was placed in the same context for 32 minutes without presentation of the tone. Twenty-four hours after extinction training or context exposure, mice were placed back in the conditioning context and presented with five non-reinforced tones to measure the amount of platform-mediated avoidance, latency to the platform, and tone-induced freezing. Freezing measurements did not distinguish between on-platform and off-platform freezing.

### 2.3 Exclusion Criteria, Behavioral, and Statistical Analysis

An exclusion criterion was set for the number of shocks received. If mice received three shocks or more during the third conditioning session, they were excluded from the remainder of the experiment. After the exclusion criterion was applied, the remaining mice were matched on the percent time on the platform before undergoing extinction. Mice were scored on the first through third training days for acquisition in the PMA paradigm and during the test session. Two experimenters scored every mouse for the total time spent on the platform during each 30-second tone presentation. Freezing behavior was assessed using FreezeFrame5 software (Actimetrics). Darting was assessed by exporting videos from FreezeFrame5 and converting them to mpeg files. Videos were analyzed using ANY-Maze (Stoleting.Co) for average speed and maximum speed. A cutoff of 23.5cm/s was used as a criterion for darting (Gruene, et al., 2015). To compare avoidance data before and after extinction training, we calculated the percentage of time spent on the platform during all five tones on the third day of training. We then compared those percentages to the average percent time on the platform during all five tones in the post-extinction test. All statistical analyses were conducted using GraphPad Prism statistical software (GraphPad Prism 10). Unpaired t-tests, and repeated measures two-way or three-way ANOVAs were used. Significant main effects were followed up with Tukey’s or Sidak’s post-hoc analyses where appropriate. All data were graphed using the standard error of the mean (SEM). Effect size and statistical power were calculated using G*Power 3.1. Please refer to tables 1 and 2 for all statistical details.

**Table 1:**
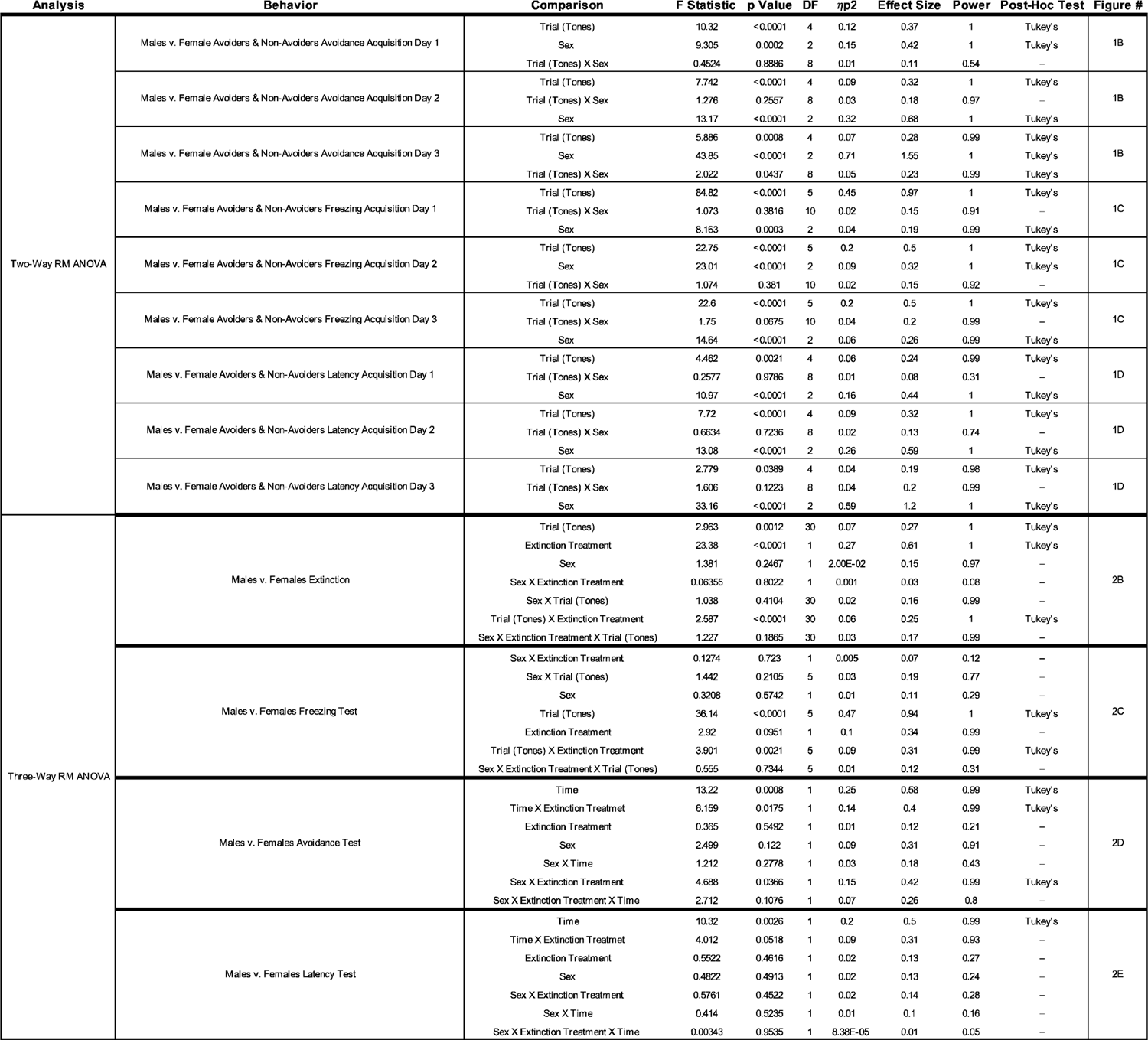
Sex differences in PMA Statistical Summary.

**Table 2:**
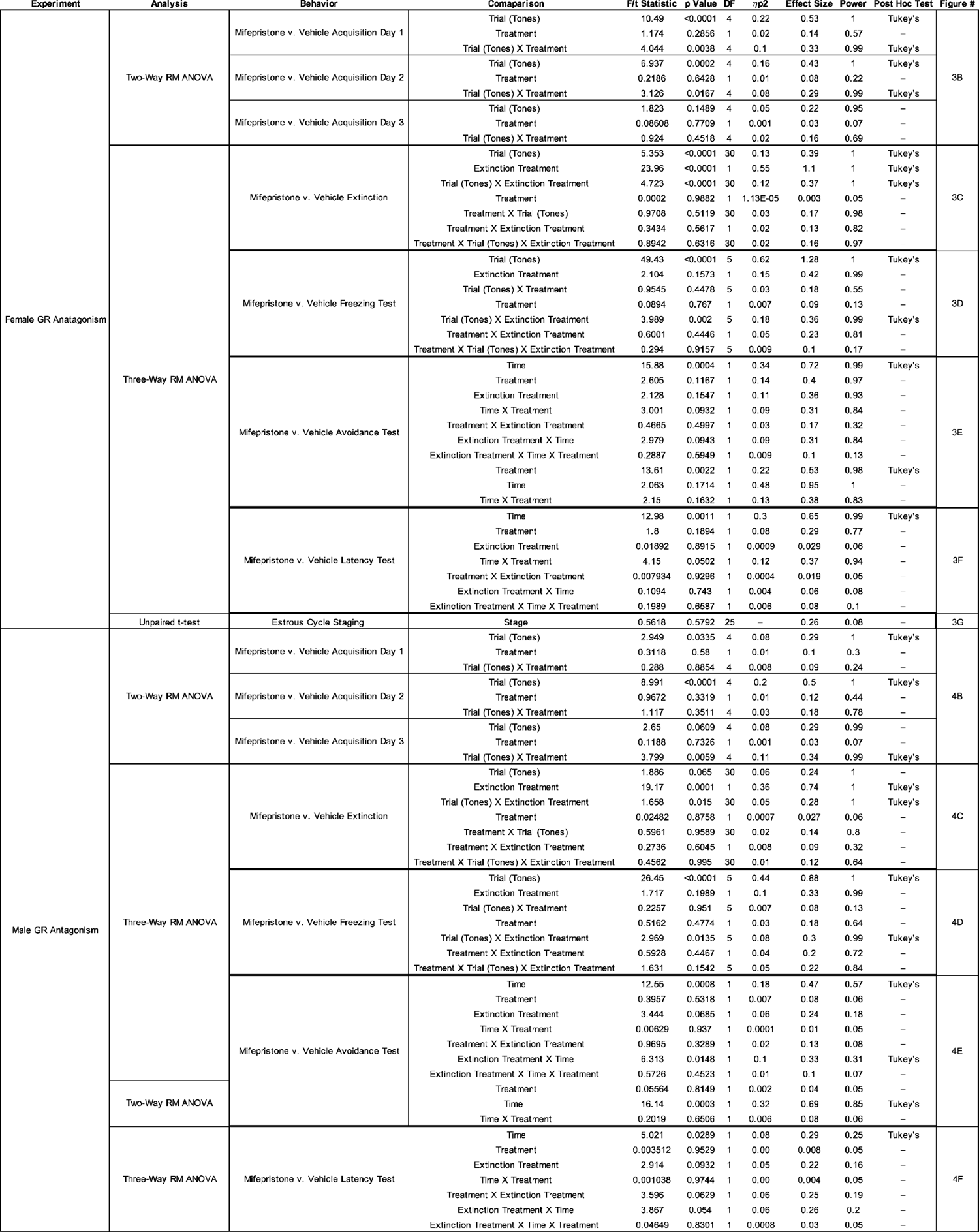
GR Antagonism Statistical Summary.

### 2.4 Drug Preparation

RU486 (mifepristone), a glucocorticoid receptor antagonist, was used to assess their role in avoidance and freezing. Mifepristone (Hello Bio) was dissolved in 10% ethanol and 90% peanut oil (Sigma) for a final concentration of 25mg/kg (Donley, et al., 2005; Okamoto, et al., 2015) set to a volume of 0.1mL per 10 grams of body weight. The vehicle solution was prepared identically, with the absence of the drug. Mifepristone was delivered via subcutaneous injection one hour before training.

### 2.5 Estrous Cycle Measurement

Female mice underwent vaginal lavage after avoidance testing in the Mifepristone experiment. Autoclaved water was gently inserted into the vagina via a sterile pipette tip attached to a latex bulb (McLean, et al., 2012), and samples were placed onto glass slides. Following sample collection, slides were left to air dry and then were dipped in 0.1% cresyl violet solution and rinsed in deionized water. Slides were examined using light microscopy to assess the cycle stage. The stage was determined by the presence of leukocytes, nucleated epithelial cells, and/or cornified epithelial cells. Saturation of the three cell types was scored and quantified using the +/- system where - = none, + = low, ++ = moderate, and +++ = high (Cora, et al., 2015).

## 3.0 Results

### 3.1 Female mice display differences in platform-mediated avoidance learning and a more persistent avoidance after extinction

For details on all statistical analyses, please see Table 1. Twenty-eight male and 50 female mice were initially run in a three-day training procedure. We observed similar avoidance using this procedure compared to studies using a 10-day procedure in rats (Bravo-Rivera, et al., 2014; Martinez-Rivera, et al., 2020; 2022). During avoidance training, we discovered that approximately 48% of the females qualified for exclusion during the third training session (3 or more shocks on the third training day) (Figure 1B). After exclusion, 19 male and 26 female mice remained in the study. We found that male and female mice that sufficiently acquired platform-mediated avoidance spent significantly more time on the platform compared to non-avoiding females by the end of training [F (2, 74) = 43.85, p < 0.0001], and during the first [F (2, 75) = 9.305, p = 0.0002] and second sessions [F (2, 75) = 13.17, p < 0.0001]. We also assessed the freezing during the tone presentations across groups (Figure 1C) and found a significant group (males, female avoiders, female non-avoiders) difference on the first day [F (2, 450) = 8.163, p = 0.0003]. We also saw a group difference on days two [F (2, 450) = 23.01, p < 0.0001] and three [F (2, 450) = 14.64, p < 0.0001]. Moreover, we also assessed the latency to mount the platform across the training days and found a main effect of group on days one [F (2, 75) = 10.97, p < 0.0001], two [F (2, 75) = 13.08, p < 0.0001], and three [F (2, 74) = 33.16, p < 0.0001].

**Figure 1:**
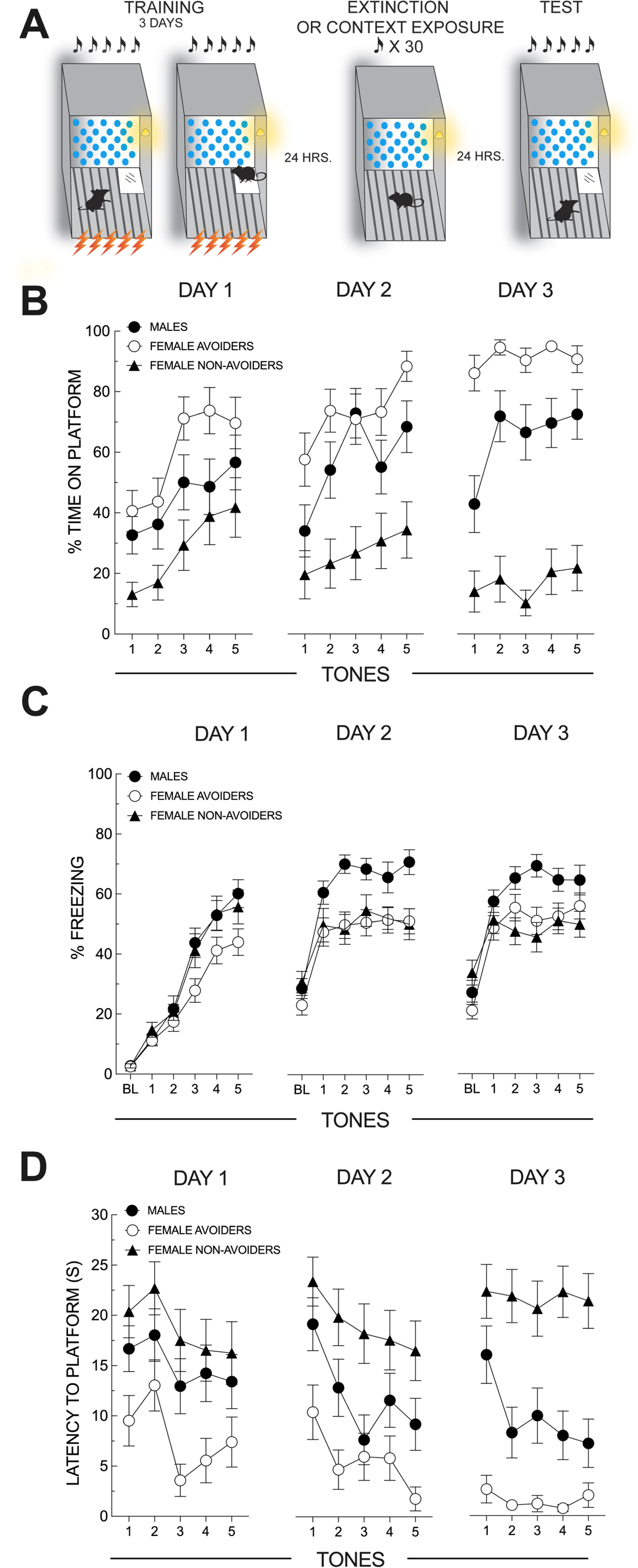
Sex differences in platform-mediated avoidance learning. **A:** Schematic of the behavioral paradigm. Mice were presented with five tone-shock pairings (75db, 6 kHz, 30s tone; 1s, 0.5mA shock). A small acrylic platform (4” x 4”) was placed in the back right corner of the chamber to provide an escape from shock for the mice. Mice were trained for three consecutive days, and time spent on the platform, latency to mount the platform, freezing, and darting were measured. **B:** Acquisition of platform-mediated avoidance; data from all mice, regardless of meeting exclusion criteria, are shown. Male mice and a subgroup of female mice (female avoiders) increased avoidance over the three-day training procedure. A separate group of female mice (female non-avoiders), however, did not increase avoidance over the training days. Female avoiders that successfully acquired platform-mediated avoidance spent significantly more time on the platform compared to non-avoiding females on the third day of training [F (2, 74) = 43.85, p < 0.0001]. **C:** Freezing during platform-mediated avoidance acquisition; data from all mice, regardless of meeting exclusion criteria, are shown. We assessed freezing during each tone presentation and found sex differences on all three training days, mainly driven by elevated freezing in male mice. **D:** Latency during acquisition; data from all mice, regardless of meeting exclusion criteria, are shown. Female avoiders and males displayed a significantly shorter latency to mount the platform during the tone compared to female non-avoiders.

Additionally, the group difference in latency to mount the platform was primarily driven by differences between female avoiders and female non-avoiders. Female non-avoiders had a higher latency to mount the platform overall than males and female avoiders (see Table 1 for individual post hoc tests). Although female non-avoiders exhibited less avoidance, they also exhibited less freezing than males but were not different from female avoiders (see above). Because avoidance was so low in female non-avoiders, we measured darting to determine what the mice were doing during the tone. We found that none of the mice showed a velocity higher than 13cm/s for females or 9.8cm/s for males, indicating a lack of darting across the training sessions. However, mice did exhibit flight responses toward the platform at tone onset if they were not on the platform, but this was not at the velocity previously established for darting. Thus, although many female mice do not acquire avoidance by day three, they do not exhibit darting as an alternative fear response strategy.

Following the third training day, mice that met the exclusion criterion were removed from the remainder of the experiment (see section 2.3). Twenty-four hours after the third training day, mice underwent extinction without the platform present or underwent context exposure also without the platform (Figure 2B). Mice that underwent extinction showed a decrease in tone freezing throughout the session, with no differences in freezing between sex (no main effect of sex [F (1, 41) = 1.381, p = 0.2467]) (Figure 2B). Mice that underwent context exposure displayed very low freezing, which was significantly lower than mice exposed to tones [F (1, 41) = 23.38, p < 0.0001], suggesting that context fear did not contribute substantially to tone-induced freezing.

**Figure 2:**
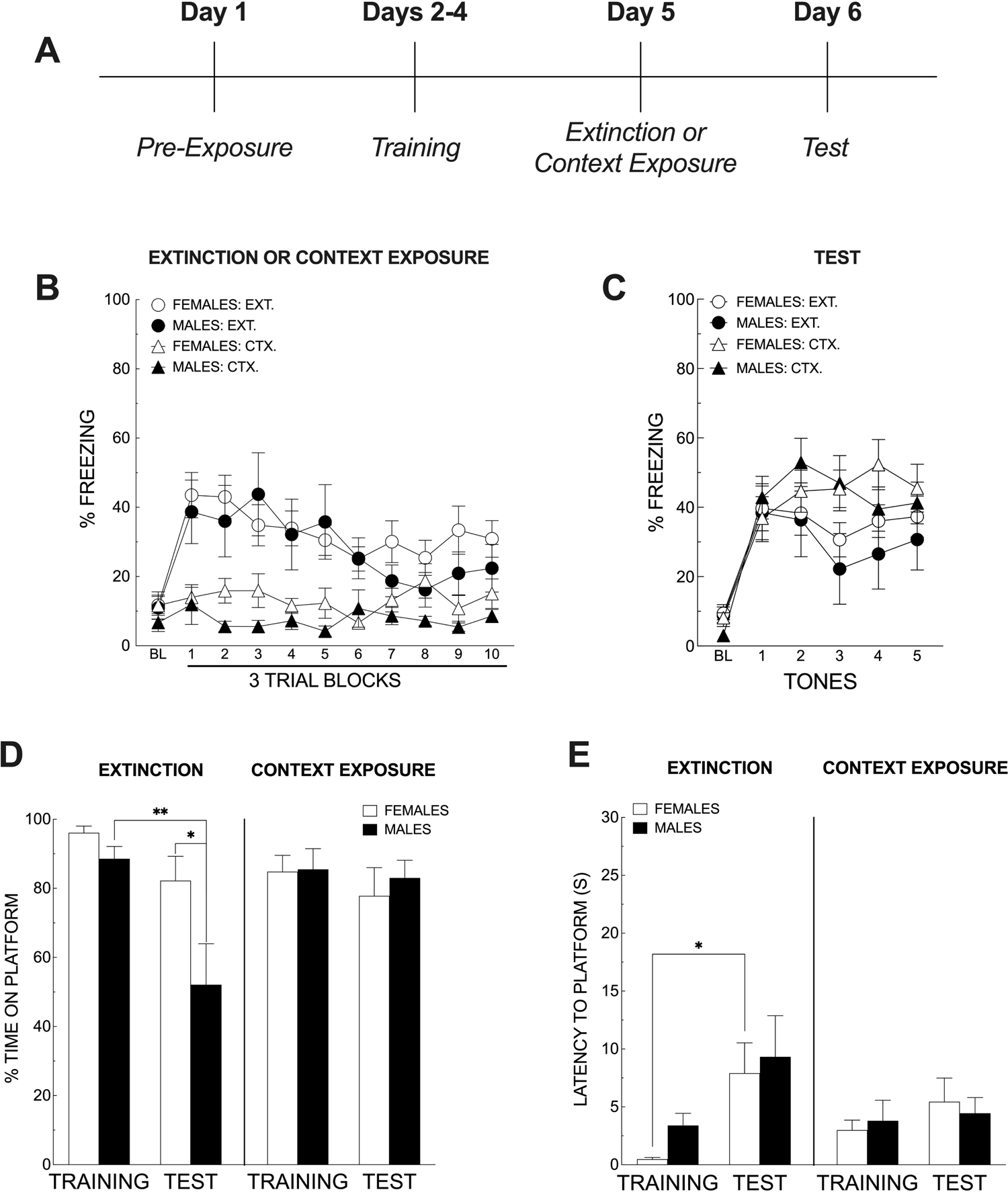
Female mice show persistent avoidance after extinction. **A:** Timeline of experiment. **B:** Male and female mice underwent a 30-tone extinction procedure with no platform present or context exposure for 32 minutes. There were no sex differences in freezing during extinction. **C:** Percent freezing to the tone during the post-extinction avoidance test (5 tone presentations). There was no significant effect of sex on freezing during the test session [F (1, 41) = 0.3208, p = 0.5742]. **D:** Pre-and Post-extinction avoidance as measured by the percent time spent on the platform only for mice that passed exclusion criteria on day three of acquisition. Data are the average time on the platform across all five tones on the last day of training compared to the average time on the platform across all five tones during the post-extinction test. Male mice in extinction conditions significantly reduced the percentage time spent on the platform (p = 0.0047). Female mice that underwent extinction did not show significant reductions in avoidance (p = 0.5785), suggesting persistent avoidance. **E**: Pre- and post-extinction latency to mount the platform compared to latency to mount the platform during the post-extinction test. There were no significant differences between males and females for the latency to mount the platform during the tone presentations. However, female mice that underwent extinction significantly increased latency to mount the platform (p = 0.0289).

One day after extinction or context exposure, mice underwent a 5-tone test session to assess freezing and avoidance. For tone freezing, there was no significant effect of sex [F (1, 41) = 0.3208, p = 0.5742], indicating that differences in persistent fear responses are exclusive to avoidance (Figure 2C). For avoidance, mice were compared using the average percent time on the platform across all five tones presented during their last day of training compared to the average percent time on the platform across all five tones presented during the 5-tone test session that occurred after extinction or context exposure. Only mice that did not meet the exclusion criterion were used for this analysis. This enabled us to assess pre-versus post-extinction avoidance (Figure 2D). We found a significant sex X extinction treatment interaction [F (1, 39) = 4.688, p = 0.0366]. Moreover, male mice that underwent extinction showed a significant reduction in platform-mediated avoidance from their training levels (p = 0.0047). In contrast, female mice that underwent extinction were no different from their training avoidance (p = 0.5785). Additionally, we found that males and females during the test were significantly different (p = 0.0307). Overall, this suggests that females display persistent avoidance even after extinction (Figure 2D) despite some reduction in avoidance. We also measured the latency to the platform (Figure 2E) and found an overall significant increase in the latency between training and test [F (1, 41) = 10.32, p = 0.0026]. Additionally, we saw that female mice showed a significant increase in latency to mount the platform following extinction (p = 0.0289). This suggests that although female mice show persistent avoidance, extinction increased their latency to move to the platform. The results reveal a significant sex difference in the acquisition of platform-mediated avoidance, where many females did not acquire avoidance based on our exclusion criterion (non-avoiders) and differences in post-extinction avoidance. Females did not extinguish platform mediated avoidance despite undergoing the same extinction training as males.

### 3.2 Mifepristone augments extinction for avoidance in female mice

Fifty-two female mice were initially run in the three-day training procedure. After exclusion, 39 mice remained in the study. However, during the test, 4 mice were not recorded due to a technical error. Therefore, the test data only contained an N of 35. Female mice received subcutaneous injections of mifepristone or vehicle one hour before each of the three training sessions (Figure 3A). Among the females that met the acquisition criterion, we found a significant main effect of tone presentations across the first two training sessions (Table 2).

**Figure 3:**
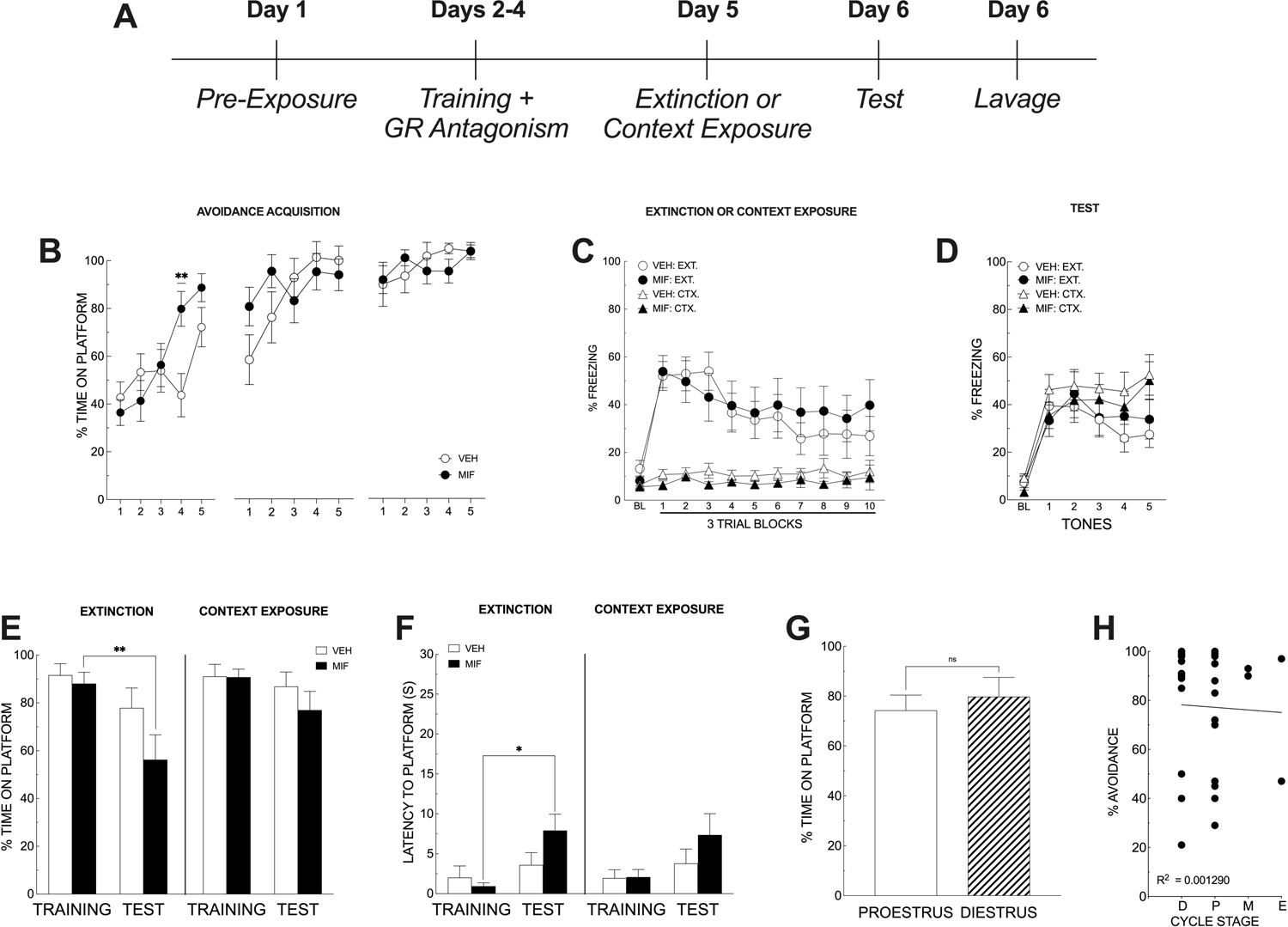
Mifepristone treatment reduces persistent avoidance in female mice. **A:** Experimental timeline. **B:** Acquisition of PMA in female mice treated with mifepristone or vehicle during the three-day training procedure. Female mice increased their avoidance across the training days. There was no overall effect of mifepristone on avoidance learning across the three-day training procedure. **C:** Twenty-four hours after the last training session, female mice underwent a 30-tone extinction or context exposure for 32 minutes. Mice exposed to extinction reduced freezing across the extinction session (3-trial blocks), whereas context-exposed mice showed little context freezing. **D:** Percent freezing during the post-extinction avoidance test (5 tone presentations). Mifepristone had no effect on freezing during the post-extinction test session. **E:** Pre- and Post-extinction avoidance as measured by the percent time spent on the platform only for mice that passed exclusion criteria on day three of acquisition. Data are the average time on the platform across all five tones on the last day of training compared to the average time on the platform across all five tones during the post-extinction test. Mifepristone-treated female mice showed significantly reduced post-extinction avoidance compared to pre-extinction avoidance (p = 0.0065). There was no significant reduction in avoidance observed in vehicle-treated females, suggesting that mifepristone treatment during avoidance learning promotes more effective extinction **F:** Pre- and post-extinction latency to mount the platform compared to latency to mount the platform during the post-extinction test. Mifepristone-treated females showed a significant increase in their latency to mount the platform after extinction compared to their pre-extinction latency (p = 0.0475). **G:** There was no significant difference in avoidance between females in proestrus and those in diestrus [t (25) = 0.5618, p = 0.5792]. **H:** Regression analysis of cycle stage and avoidance, regardless of treatment. There was no significant relationship between stage and avoidance (D = Diestrus; P = Proestrus; M = Metestrus; E = Estrus) [R^2^ = 0.001290, p = 0.8479].

Additionally, there was a significant tone X treatment interaction on day one [F (4, 148) = 4.044, p = 0.0038] and day two [F (4, 148) = 3.126, p = 0.0167]. Tukey’s post hoc analysis of day one revealed a significant increase in percent time on platform for the mifepristone group on the fourth tone compared to vehicle treated mice (p = 0.0037). (Figure 3B). However, we did not find a main effect of treatment on any of the days (Figure 3B). Mice were matched on their avoidance based on the last day of training and placed into extinction or context exposure groups (Figure 3C). Mice that underwent extinction showed reductions in tone freezing across the session [F (3, 35) = 5.353, p < 0.0001], but there was no effect of treatment on freezing in the extinction and context exposure groups [F (1, 35) = 0.0002, p = 0.9882] (Figure 3C). Twenty-four hours later, mice were returned to the chamber and underwent a 5-tone test. Freezing was not different between mifepristone-treated and vehicle-treated mice [F (1, 30) = 0.0894, p = 0.7670], (Figure 3D). However, mice that had received mifepristone and underwent extinction showed a significant reduction in the percent time on the platform compared to their training (pre-extinction) time (p = 0.0065; 3-Way, p = 0.0019; 2-Way). There was no difference in platform mediated avoidance in vehicle-treated females that underwent extinction (p = 0.7161), suggesting that extinction alone was ineffective at reducing avoidance in females (Figure 3E). We also found that latency to the platform was significantly longer in those mice that underwent extinction and were treated with mifepristone (p = 0.0475) (Figure 3F).

Mifepristone thus enhanced the effectiveness of extinction training to reduce persistent avoidance in females. We then analyzed the estrous stage of all mice, regardless of treatment conditions. We grouped them into proestrus (highest levels of estradiol) or diestrus (lowest levels of estradiol). There was no significant difference in avoidance across cycle stage [t (25) = 0.5618, p = 0.5792], regardless of treatment condition (Figure 3G). We also analyzed the data using a simple linear regression (D = Diestrus; P = Proestrus; M = Metestrus; E = Estrus) and found no significant relationship between cycle stage and levels of platform mediated avoidance [R^2^ = 0.001290, p = 0.8479] (Figure 3H). The results suggest that GRs during avoidance learning may be responsible, in part, for persistent avoidance in females, and blocking their activity during acquisition augmented the effectiveness of extinction on avoidance, and the effects we see are not dependent on estrous cycle stage.

### 3.3 Mifepristone doesn’t augment extinction for avoidance in male mice

Forty male mice were initially run in the three-day training procedure. After exclusion, 38 mice remained in the study. However, during the third training day, 4 mice were not included due to a software error and data not being recorded. Therefore, their data was not in the statistical analysis, leaving 34 mice in the analysis. Male mice received mifepristone or vehicle injections one hour before each of the three training sessions (Figure 4A). Across days, mice increased their levels of avoidance (Figure 4B). We found a significant effect of tone presentations in the first two training sessions (Table 2). On the third day, we saw a significant tone X treatment interaction [F (4, 128) = 3.799, p = 0.0059]. However, there was no main effect of treatment across any of the training sessions; day 1 [F (1, 36) = 0.3118, p = 0.5800], day 2 [F (1, 36) = 0.9672, p = 0.3319], and day 3 [F (1, 32) = 0.1188, p = 0.7326). Mice were matched on avoidance as described above and, twenty-four hours after the last day of training were placed into the conditioning chamber to undergo extinction or context exposure (Figure 4C). There was a main effect of extinction and a significant tone X extinction interaction (Table 2). Twenty-four hours after extinction or context exposure, mice were tested for freezing, avoidance, and latency to mount the platform. Freezing during the test was not different between vehicle and mifepristone-treated mice [F (1, 34) = 0.5162, p = 0.4774] (Figure 4D). During the avoidance test, we found a main effect of time [F (1, 58) = 12.55, p = 0.0008] and a significant time X extinction treatment interaction [F (1, 58) = 6.313, p = 0.0148]. Male mice that underwent extinction significantly reduced their percent time on the platform regardless of drug treatment (p = 0.0171 for mifepristone; p = 0.0032 for vehicle) (Figure 4E). Finally, we assessed latency to mount the platform (Figure 4F) and overall, a significant increase in the latency to mount the platform during the test compared to training [F (1, 58) = 5.021, p = 0.0289]. However, we did not find any significant post-hoc comparisons for the groups. Overall, this suggests that mifepristone treatment in males during avoidance learning does not reduce avoidance more than extinction alone and has no effect on freezing or latency to approach the platform.

**Figure 4:**
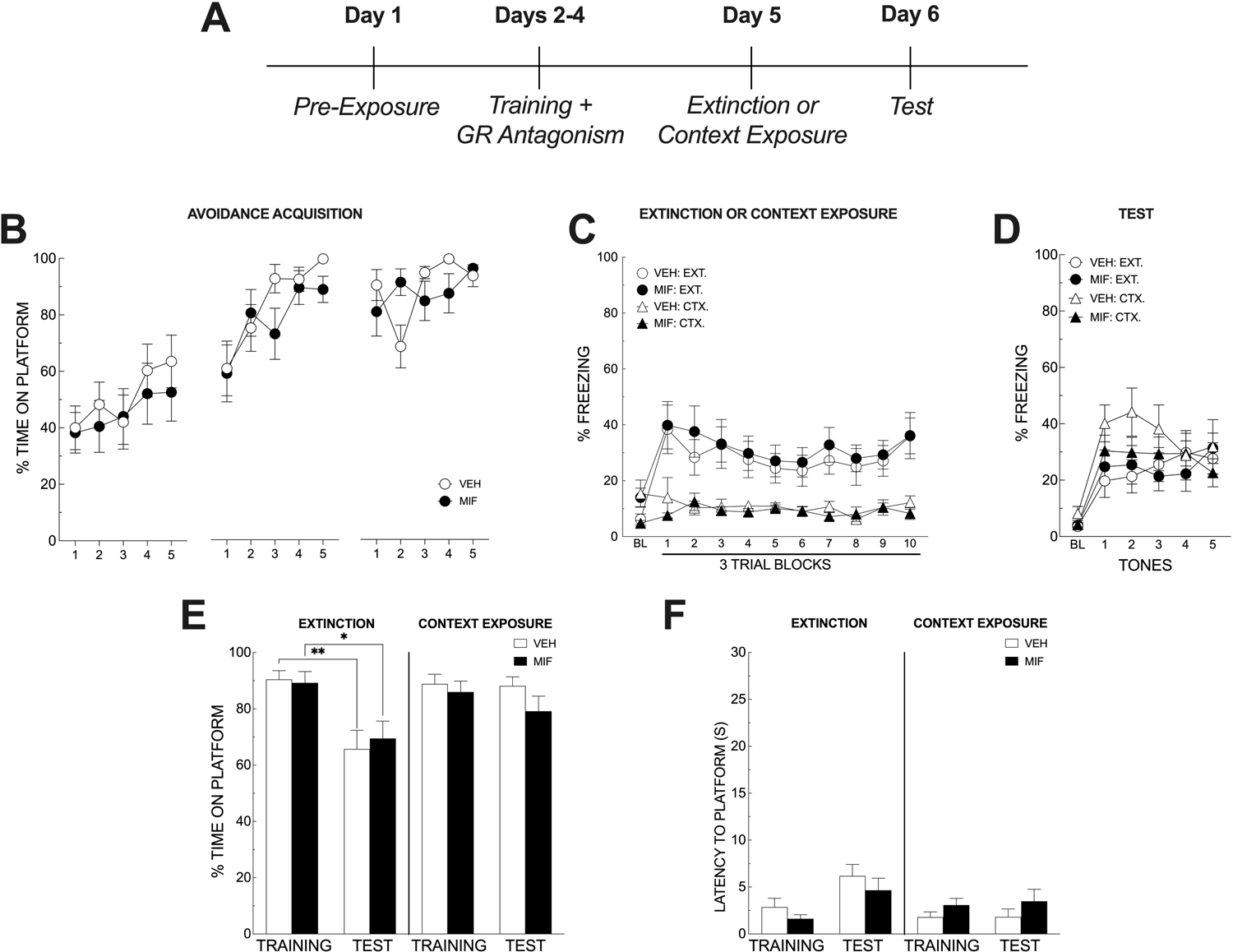
Mifepristone treatment does not further reduce avoidance in male mice. **A:** Experimental timeline. **B:** Male mice increased their time spent on the platform across the three training sessions, but there was no overall effect of mifepristone on avoidance across the three training sessions. **C:** Extinction or context exposure treatment following the 3-day training sessions. There were no significant effects of mifepristone treatment on tone freezing during extinction or in the context exposure groups. **D:** Tone freezing during the post-extinction test session. Mifepristone also had no effect on tone-induced freezing during the post-extinction test session. **E:** Pre- and Post-extinction avoidance as measured by the percent time spent on the platform only for mice that passed exclusion criteria on day three of acquisition. Data are the average time on the platform across all five tones on the last day of training compared to the average time on the platform across all five tones during the post-extinction test. Mifepristone provided no additional reduction in avoidance beyond extinction and vehicle treatment. **F:** Pre- and post-extinction latency to mount the platform compared to latency to mount the platform during the post-extinction test. Mifepristone had no effect on latency to mount the platform during the test session.

## 4.0 Discussion

The current study is the first to examine platform-mediated avoidance in female mice and revealed a striking sex difference in the acquisition and post-extinction avoidance behavior. Notably, male and female rats have been previously used in PMA but displayed similar behavioral phenotypes (Bravo-Rivera, et al., 2021). Across all experiments, in our three-day training procedure, 83.8% of male mice avoided shock on the third day and met the training criterion to proceed with the extinction phase of the experiment. Males also displayed a progressive acquisition curve during the training trials (Figure 1B). Of the female mice that met the training criterion on day three (across all experiments, 63.7%), their percent avoidance remained similar to males over the first two days, then was elevated on day three (Figure 1B). The remainder of the female mice did not efficiently avoid the shocks on the second and third training days (i.e., they received three shocks on day three), suggesting male mice employ a single avoidance strategy to deal with cues predicting threat. However, it should be noted that freezing was also elevated due to the lack of an appetitive task like that in Bravo-Rivera, et al., 2014. Thus, avoidance is not the only behavior displayed during CS presentations; freezing responses are also engaged.

In contrast, female mice may engage in divergent strategies that include acquiring avoidance and non-avoidance of predictive aversive cues. Another possibility is that this subgroup of females acquire avoidance more slowly, an effect that could be revealed by allowing them to proceed with a prolonged training procedure. One final possibility in the non-avoiding females is that their engaging freezing inhibits their ability to engage in an avoidance response (Lazaro-Munoz, et al.,2010). The estrous cycle stage can affect the acquisition of aversive learning (Trask, et al., 2020; Carvalho, et al., 2021). Therefore, we cannot rule out the effects of cycle stage on avoidance learning during the acquisition phase of the present study. A similar sex difference in shock avoidance has been reported using nose-poke responding to avoid shock (Kutlu et al., 2020). How these divergent behaviors evolved, and their underlying mechanism is currently unclear, although we do not think it is estrous cycle-dependent based on the multi-day procedure and our cycle stage data from the mifepristone experiment (see discussion below). In addition, across all experiments, 16.2% of males did not meet the criterion on day three, suggesting a smaller percentage engaged in freezing over avoidance, but this would not have been influenced by cycle stage. In the post-extinction avoidance test, males significantly reduced their time avoiding (less time spent on the platform) after the three-day training procedure. However, females that acquired avoidance during the same training procedure did not reduce their time on the platform after extinction (Figure 2D). Thus, female mice that acquire avoidance display more persistent avoidance behavior than males despite similar training and extinction conditions. This could be partly explained by differences in avoidance expression or could be artificial given that female avoidance is near ceiling (Figure 2D). However, avoidance during pre-extinction between males and females was not statistically different (p = 0.9913) and there was a significant reduction in avoidance observed in males.

Males and females have been reported to display separate mechanisms (behaviorally and biologically) for various learning tasks, such as fear learning, fear extinction, and fear generalization (for review, see Frick et al. 2010, 2018; Adkins et al. 2019). Several reports demonstrate that males display increased contextual fear learning compared with females (Maren et al. 1994; Markus and Zecevic 1997; Gresack et al. 2009; Mizuno and Giese 2010).

One reason for this might be that female rats are less likely to use contextual cues to recall associative learning and may rely more on other non-contextual cues for recall (Anderson and Petrovich 2015, 2018a,b). Here, we demonstrated a difference in cue-associated avoidance during acquisition and after extinction. Under the same training conditions, female mice show increased avoidance on the last day of training compared to males and display persistent avoidance after undergoing extinction. Male mice showed reduced avoidance after undergoing the same extinction procedure. Male and female mice in the context exposure groups displayed very low freezing to the context suggesting that fear of the context itself was unlikely to drive freezing or avoidance. This divergent avoidance response could be helpful in investigating mechanisms related to the sex differences observed in the rates of anxiety in humans, in which avoidance is a key symptom. However, the mechanism underlying this divergent avoidance response is currently unknown.

Unlike the procedure used here, the previous literature establishing the platform-mediated avoidance task in rats integrated an appetitive task and used training procedures that lasted 10-20 days (Bravo-Rivera, et al., 2014, Martinez-Rivera, et al., 2020; 2022). Although the rats in those experiments acquired avoidance to the same degree by the third day of training, like we observed here, freezing was consistently suppressed starting on day six. This was also when lever pressing for food reward started to return to pre-training frequency. Despite not having a motivated task to draw the mice off the platform, avoidance can be acquired at similar rates, but freezing remains comparatively high throughout our procedure. However, this makes our modified design more like one-way active avoidance but still allows a place where shock can be avoided, unlike two-way active avoidance. Interestingly, the context exposure groups displayed very low contextual freezing. We think this occurs for several reasons. First, freezing overall, even to the CS is consistently lower during platform-mediated avoidance, especially given the number of training trials compared to traditional cued fear conditioning. Second, while the lack of context freezing is unexpected, removing the platform during the extinction sessions could represent a significant enough context shift to reduce freezing. Third, freezing to the tone CS and context freezing during the extinction sessions with the platform removed is significantly lower than that observed during the last day of training with the platform present (even when mice were not receiving shocks), suggesting that the platform itself may influence freezing.

Finally, the tone is the most salient threat signal, and there are 15 training trials during acquisition. This may enhance discrimination between the context and the tone CS, facilitating reduced contextual freezing. It should be noted that we did not distinguish between on-platform and off-platform freezing. However, latencies to mount the platform were uniformly below 5 seconds before extinction training, suggesting most freezing occurred during avoidance. It will be interesting to observe if similar sex differences in acquisition and post-extinction avoidance remain when we incorporate a food reward task into our procedure for future studies.

Glucocorticoid release and glucocorticoid receptor (GR) activation have long been associated with enhanced memory for avoidance tasks, but this has mainly been investigated using one-trial inhibitory avoidance (e.g., Roozendal and McGaugh, 1997; Chen et al., 2012; Scheinman et al., 2018; Lingg et al., 2020). Platform-mediated avoidance utilizes a multi-day training procedure; thus, we were interested in determining the effects of GR blockade on the acquisition of avoidance and its effects after extinction in the absence of the platform. The GR antagonist Mifepristone (RU-486) was administered to male and female mice to assess avoidance, freezing, and latency behaviors. We wanted to determine if GR antagonism would block the persistent avoidance in females under the three-day training procedure, and how GR blockade would affect males who already show reductions following extinction. There was an effect of mifepristone on the acquisition of avoidance in females, for the first training session (Figure 3B). Previous reports have shown an effect of mifepristone on cue reconsolidation (Pitman, et al., 2011); however, our study did not see a main effect of mifepristone treatment across training sessions. We also did not observe an effect of mifepristone on freezing during extinction training within our female mice (Figure 3C) or during the test session (Figure 3D).

However, we did find that GR antagonism during acquisition paired with extinction significantly reduced avoidance responses (Figure 3E) and increased latency to mount the platform during tone presentation (Figure 3F). When assessing GR blockade in male mice, we did not see an effect of treatment across the training sessions (Figure 4B) and there was no effect on freezing during extinction (Figure 4C) or test (Figure 4D). Finally, we found that mifepresitone did not further reduce avoidance over extinction alone in males (Figure 4E). These data suggest that GR activation in females during avoidance learning partly drives persistent avoidance even after mice undergo extinction training. On the other hand, GR activation in males during avoidance learning may play a less prominent role in persistent avoidance.

These data suggest GRs may play a prominent role in regulating persistent avoidance in females, although we cannot rule out differences in GR sensitivity to mifepristone or the progesterone receptor-blocking actions of mifepristone. Blocking GRs with systemic mifepristone can inhibit negative feedback on the HPA axis (Heitzer, et al., 2007). Mifepristone treatment during avoidance training could promote an increase in plasma corticosterone that then facilitates the proper extinction of avoidance responses in female mice. Similarly, administering corticosterone after cued fear conditioning enhances extinction in male and female mice 24 hours later (Lesuis, et al., 2018). Moreover, there is also the potential for non-genomic mechanisms to occur, such as an increase in pre-synaptic glutamate or trafficking of AMPA receptors to the post-synaptic membrane (Popoli, et al., 2012; Musazzi, et al., 2010).

These non-genomic effects can allow for increased activation and the promotion of synaptic plasticity within regions outside of the HPA axis (e.g., amygdala, nucleus accumbens, prefrontal cortex) and are also known to be critical in regulating avoidance responses. Currently, we do not know if mifepristone acts within these brain regions, or its effect is primarily mediated via changes to corticosterone regulation and negative feedback of the HPA axis. It is known that the prelimbic cortex (PL) and basolateral amygdala (BLA) are critical in regulating freezing responses to tone (Sierra-Mercado, et al., 2011; Corcoran & Quirk, 2007).

Moreover, previous findings showed that inactivation of the BLA prior to avoidance testing resulted in reductions in freezing and avoidance in PMA (Bravo-Rivera, et al., 2014). When females were given mifepristone during avoidance training, they did not show differences in tone evoked freezing during testing (Figure 3D). However, they did display a reduction in avoidance responses (Figure 3E). This could be due to different strategies in fear responding, in which female mice display a preference for active avoidance over freezing (Gruene, et al., 2015; Colom-Lapetina, et al., 2019). Note, however, that a large percentage of the females in this study did not acquire avoidance according to our criterion, suggesting that adopting an avoidance strategy is not the primary response in female mice. It has also been identified that inactivation of the PL, or nucleus accumbens (NAc) during the PMA test reduces avoidance responses (Bravo-Rivera, et al., 2014). Therefore, mifepristone’s actions could act within these regions to facilitate proper behavioral responses following extinction. Additional studies are needed to elucidate these sex differences in GR regulation of avoidance.

There are sex differences in HPA axis regulation (Handa, et al., 1994; Goldstein, et al., 2010), which can depend on the estrous cycle phase. Therefore, we staged the estrous cycle in the female mifepristone experiment to assess the potential role of cycle stage on avoidance and freezing responses after GR antagonism. We did this after observing that a large percentage of females did not acquire the avoidance response in the first experiment. During diestrus, when estradiol levels are at their lowest (Ajayi & Akhigbe, 2020), baseline levels of corticosterone are low, with an optimal return to baseline after stress. However, during proestrus, when estradiol levels are high, there is an elevated baseline level of corticosterone accompanied by a delayed return to baseline after stress (Carey, et al., 1995; Viau, et al., 1991). To investigate the role of the estrous cycle in persistent avoidance in females, we staged the estrous cycle after the post-extinction avoidance test in the mifepristone experiment. Mice were staged, and we analyzed the avoidance regardless of treatment. Most females fell into diestrus or proestrus categories, but there was no statistically significant difference in avoidance between these groups of females (Figure 3G). We also ran a regression across all cycle stages to determine if cycle stage predicted avoidance. Cycle stage did not predict avoidance (Figure 3H), suggesting that differences in avoidance are unlikely to be driven by cycle stage in this experiment.

The current study is the first to show sex differences in PMA acquisition and post-extinction avoidance in mice. Male mice appear to have a single strategy for acquiring avoidance, whereas females may have different strategies that include avoidance and freezing. This highlights the importance of incorporating both sexes and multiple behavioral analyses across studies using animal models. Our findings also demonstrate a novel mechanism by which female mice display persistent avoidance compared to males with relatively few training sessions. GRs activation during learning may promote more stable avoidance learning that is resistant to a single extinction session. Future investigations regarding the mechanism through which GRs facilitate persistent fear responses across sexes will be needed to identify specific mechanisms that regulate these divergent responses. The current findings also highlight the therapeutic potential of GR antagonism for disorders involving persistent avoidance.

## Acknowledgements

We acknowledge the assistance of the animal care staff at the Psychological Sciences Department of Kent State University and the University of South Carolina School of Medicine. Additionally, we acknowledge Kaitlin Martin, Danielle Rodgers, and James Hoyt for their assistance in behavioral scoring.

